# Exploring sex differences in the prevention of ethanol drinking by naltrexone in dependent rats during abstinence

**DOI:** 10.1101/310334

**Authors:** A Matzeu, L Terenius, R Martin-Fardon

**Affiliations:** Department of Neuroscience, The Scripps Research Institute, La Jolla, CA, USA; Karolinska Institute, Clinical Neuroscience, Experimental Addiction Research, Stockholm, Sweden.

**Author notes:** Corresponding Author: Dr. A. Matzeu, Department of Neuroscience, The Scripps Research Institute, 10550 North Torrey Pines Road, SP30-2003, La Jolla, CA 92037, USA.

**Keywords:** alcohol, ethanol, intake, abstinence, naltrexone, dependence

## Abstract

**Background:** Despite considerable efforts, few drugs are available for the treatment of alcohol (ethanol [EtOH]) use disorders (AUDs). Ethanol directly or indirectly modulates several aspects of the central nervous system, including neurotransmitter/neuromodulator systems. Relapse vulnerability is a challenge for the treatment of EtOH addiction. Ethanol withdrawal symptoms create motivational states that lead to compulsive EtOH drinking and relapse even after long periods of abstinence. Among the therapeutics to treat AUDs, naltrexone (NTX) is a pharmacological treatment for relapse. The goal of the present study was to evaluate the effect of NTX on EtOH drinking in EtOH-dependent male and female rats during abstinence.

**Methods:** Wistar rats (males and females) were first trained to orally self-administer 10% EtOH. Half of them were then made dependent by chronic intermittent EtOH (CIE) vapor exposure, and the other half were exposed to air. Using this model, rats exhibit somatic and motivational signs of withdrawal. At the end of EtOH vapor (or air) exposure, the rats were tested for the effects of NTX (10 mg/kg, p.o.) on EtOH self-administration at three abstinence time points: acute abstinence (8 h, A-Abst), late abstinence (2 weeks, L-Abst), and protracted abstinence (6 weeks, P-Abst).

**Results:** NTX decreased EtOH intake in nondependent rats, regardless of sex and abstinence time point. In post-dependent rats, the effects of NTX improved with a longer abstinence time (i.e., L-Abst and P-Abst) in males, whereas it similarly reduced EtOH drinking in females at all abstinence points.

**Conclusions:** The data suggest that the therapeutic efficacy of NTX depends on the time of intervention during abstinence and sex. The data further suggest that EtOH dependence induces different neuroadaptations in male and female rats, reflected by differential effects of NTX. The results underscore the significance of considering the duration of EtOH abstinence and sex for the development of pharmacotherapeutic treatments for AUD.

## Introduction

Ethanol (EtOH) produces its effects on the central nervous system through several mechanisms, one of which is alteration of the endogenous opioid system. In the case of the endogenous opioid system and its receptor subtypes (μ, δ, κ) that are selective for the three main classes of endogenous opioids (β-endorphin, enkephalins, and dynorphins, respectively), acute EtOH stimulates the release of β-endorphin, enkephalins, and dynorphin in humans and rats (Gianoulakis et al., 1996b; Marinelli et al., 2003; Marinelli et al., 2004; Marinelli et al., 2005; Marinelli et al., 2006; Dai et al., 2005; Popp and Erickson, 1998; Rasmussen et al., 1998).

Naltrexone (NTX) is an approved medication for the treatment of alcoholism, based on its efficacy in reducing the craving for and consumption of EtOH (Volpicelli et al., 1992). It is a nonspecific opioid receptor antagonist. In nondependent subjects, blockade of the opioid receptors for these endogenous ligands by NTX suppresses EtOH consumption (Gonzales and Weiss, 1998; Coonfield et al., 2002; Shoemaker et al., 2002; Stromberg, 2004), and this beneficial effect is blunted in dependent subjects (Ji et al., 2008; Sabino et al., 2006; Sabino et al., 2013; Walker and Koob, 2008). Although clinical trials have demonstrated the efficacy of NTX in reducing the risk of relapse in heavy drinkers, many patients experience no benefit of treatment (Krystal et al., 2001). This suggests that several factors influence the success of treatment (e.g., the time of intervention during abstinence, gender, and changes in the endogenous opioid system). Numerous investigations have been sought to characterize biochemical modifications of the endogenous opioid system that are closely associated with the incidence of alcoholism or alcohol use disorder (AUD). For example, both mice and humans with a high risk of alcoholism present greater hypothalamic β-endorphin activity (Gianoulakis et al., 1996a). Repeated EtOH administration induces both short- and long-term alterations of opioid levels in brain regions that are associated with motivation and reward (e.g., nucleus accumbens; Lindholm et al., 2000). In rodents, chronic EtOH intake and EtOH withdrawal have been shown to induce a wide range of perturbations of the opioid system, suggesting a putative role for the endogenous opioid system in the effects of EtOH, such as a decrease in μ-opioid receptor expression in the nucleus accumbens and dorsal striatum (Turchan et al., 1999), an increase in prodynorphin mRNA levels in the nucleus accumbens (Przewlocka et al., 1997), a decrease in κ-opioid receptor mRNA expression in the ventral tegmental area and nucleus accumbens (Rosin et al., 1999), an increase in preproenkephalin mRNA expression in the amygdala, and a decrease in preproenkephalin mRNA expression in the nucleus accumbens (Cowen and Lawrence, 2001).

Resumption (i.e., relapse) to excessive EtOH drinking even long after physical signs of withdrawal have dissipated is a challenge for the successful treatment of AUD. Although NTX is an approved treatment for AUD, unclear is why the efficacy of treatment varies between subjects. Therefore, to gain better knowledge on the efficacy of NTX to prevent the recurrence of EtOH drinking during abstinence, this study systematically tested the effects of NTX at a dose that was reported in previous studies to be efficient in reducing EtOH consumption (Stromberg et al., 2002) at different time points during abstinence in male and female rats.

## Materials and Methods

### Rats

Forty-eight Wistar rats (24 males and 24 females; Charles River, Wilmington, MA, USA), weighing 150-175 g upon arrival, were housed two per cage in a temperature- and humidity-controlled vivarium on a reverse 12 h/12 h light/dark cycle with *ad libitum* access to food and water. The animals were given at least 1 week to acclimate to the housing conditions and handling before testing. All of the procedures were conducted in strict adherence to the National Institutes of Health *Guide for the Care and Use of Laboratory Animals* and were approved by the Institutional Animal Care and Use Committee of The Scripps Research Institute.

### Ethanol self-administration training (Fig. 1)

The EtOH self-administration training procedure was a modification of the early studies by Wise (1973) and revised by Simms et al. (2008). The procedure did not require any fading (saccharin or sucrose) procedure to initiate voluntary EtOH drinking. On the Monday (Day 1) following the end of the housing acclimatization period, the rats were single-housed and presented with two bottles: one bottle with H2O and one bottle with 10% (w/v) EtOH (prepared in tap water from 95% w/v EtOH) for 3 days (Monday, Tuesday, Wednesday; Days 1-3). On Thursday and Friday (Days 4 and 5), the rats were grouped-housed (two per cage) and given H2O only in their home cage. On Saturday (Day 6), the rats were given access to EtOH self-administration for 12 h in standard operant conditioning chambers (29 cm × 24 cm × 19.5 cm; Med Associates, St. Albans, VT, USA) on a fixed-ratio 1 schedule of reinforcement, in which each response at the right active lever (i.e., the only lever available at that time) resulted in the delivery of 0.1 ml of fluid and brief illumination of a cue light (0.5 s) above the lever. Food and water were available in the operant chambers. Following the 12 h session, the rats were returned to their home cage. On Sunday (Day 7), they were left undisturbed. On Monday and Tuesday (Days 8 and 9), the rats were given access to EtOH self-administration for 2 h/day and on Wednesday and Thursday (Day 10 and 11) for 1 h/day. Starting on Friday (Day 12) and for the rest of the self-administration days, the rats underwent 30 min self-administration sessions with the introduction of a second (left) inactive lever at which responses were recorded but had no programmed consequences.

### Chronic intermittent EtOH vapor exposure (Fig. 1)

At the end of EtOH self-administration training, two rats (one male and one female) did not acquire EtOH self-administration, thus reducing the number of animals to 46. After 21 sessions of EtOH self-administration, half of the rats (*n* = 22; 11 males and 11 females) were then made dependent (EtOH post-dependent [EtOHPD]) by chronic intermittent EtOH (CIE) vapor exposure. The other half (*n* = 24; 12 males and 12 females) comprised the nondependent group (EtOHND) that was exposed only to air. During 6-week dependence induction, the rats underwent cycles of 14 h ON (blood alcohol levels [BALs] during vapor exposure ranged between 150 and 250 mg%, measured with a blood analyzer [GC-headspace, Agilent Technologies, Santa Clara, CA, USA]) and 10 h OFF and were left undisturbed for 3 weeks except to control their BALs (measured during the last 15 min of vapor exposure) and be scored for somatic withdrawal signs (at 8 h of abstinence) once per week. Behavioral signs of withdrawal were measured using a rating scale that was adapted from an original study by Macey et al. (1996) and included ventromedial limb retraction, vocalization (i.e., irritability to touch), tail rigidity, abnormal gait, and body tremors. Each sign was given a score of 0-2, based on the following severity scale: 0 = no sign, 1 = moderate, and 2 = severe. The sum of the 5 scores (0-10) was used as a quantitative measure of withdrawal severity and to confirm dependence. In this model, rats exhibit somatic and motivational signs of withdrawal (Vendruscolo and Roberts, 2014). Starting at the beginning of the fourth week of CIE exposure, the rats were subjected to 30 min FR1 EtOH self-administration sessions when acute abstinence occurred (i.e., 8 h after the vapor was turned off when brain and blood alcohol levels were negligible), three times per week (Monday, Wednesday, and Friday), in which a gradual increase in EtOH intake should be measured (Vendruscolo and Roberts, 2014). The air-exposed rats underwent the same procedure.

### Effects of naltrexone on EtOH self-administration during abstinence (Fig. 1)

**Figure 1.**
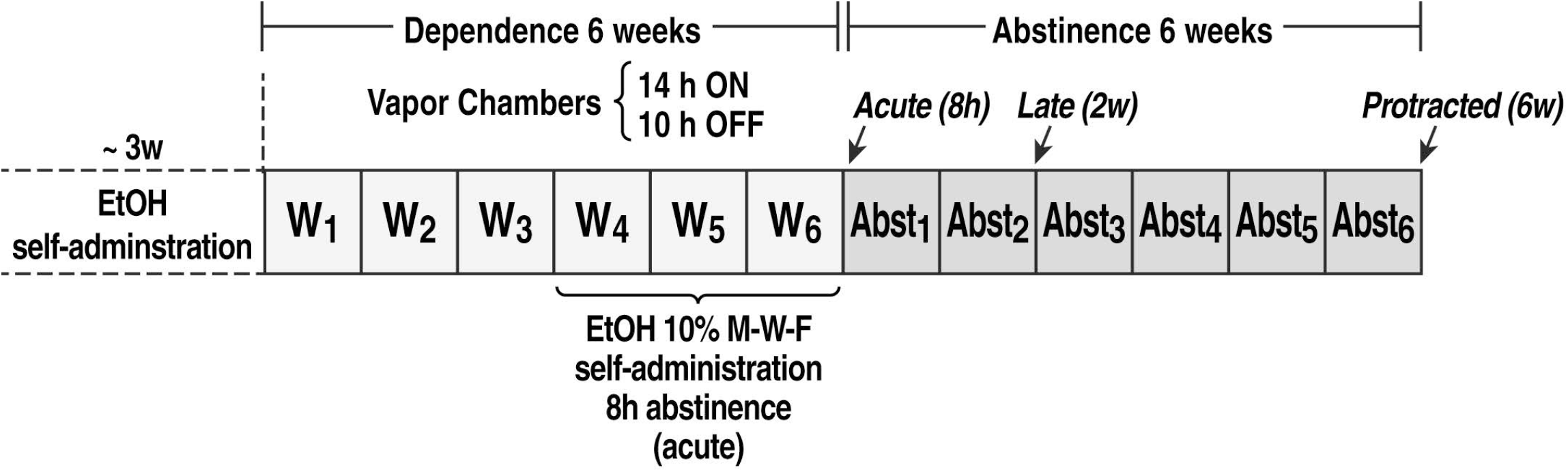
Behavioral procedure. W, week.

At the end of week 6 of CIE or air exposure, the animals began the abstinence phase and were tested for the effect of naltrexone (NTX; naltrexone hydrochloride, Sigma) on EtOH self-administration at three abstinence points: acute abstinence (A-Abst, 8 h), late abstinence (L-Abst, 2 weeks), and protracted abstinence (P-Abst, 6 weeks). At each time point of abstinence, the rats received oral NTX (10 mg/kg in a volume of 3 ml/kg; Stromberg et al., 2002; *n* = 6) or vehicle (water, *n* = 5/6 animals) administration. Sixty minutes later, they were placed in the operant chambers and allowed to self-administer 10% EtOH on an FR1 schedule for 30 min. NTX (or vehicle) was administered according to a within-subjects design, such that the same group of rats received three NTX (or vehicle) treatments, one at A-Abst, one at L-Abst, and one at P-Abst.

### Statistical analysis

One-, two-, or three-way analysis of variance (ANOVA) was used to analyze differences in EtOH intake during training and CIE exposure and after NTX treatment. Separate one-way ANOVAs or *t*-tests were used to analyze EtOH intake and BALs during CIE exposure and after NTX treatment. Withdrawal signs were analyzed using the nonparametric Kruskal-Wallis test. The results are expressed as mean ± SEM. Significant effects in the ANOVAs were followed by appropriate *post hoc* tests. Differences were considered significant at *p* < 0.05.

## Results

Two animals were excluded from the study, one in the male EtOHPD group and one in the female EtOHPD group, because they did not acquire EtOH self-administration, thus reducing the number of animals to 46 (*n* = 12 males EtOH_ND_ and *n* = 11 males EtOH_PD_ during training and CIE exposure; *n* = 6 males EtOH_ND_ treated with NTX or vehicle, *n* = 6 males EtOH_PD_ treated with NTX, and *n* = 5 males EtOH_PD_ treated with vehicle; *n* = 12 females EtOH_ND_ and *n* = 11 females EtOHPD during training and CIE procedure; *n* = 6 females EtOH_ND_ treated with NTX or vehicle, *n* = 6 females EtOH_PD_ treated with NTX, and *n* = 5 females EtOH_PD_ treated with vehicle).

### EtOH self-administration training

During the last 5 days of self-administration training (Fig. 2), both groups of animals (males and females) presented stable and comparable EtOH intake in a 30-min daily session (two-way ANOVA: sex [male *vs*. female], *F*_1,44_ = 0.1, *p* > 0.05; time [days of self-administration], *F*_4,176_ = 1.3, *p* > 0.05; sex × time interaction, *F*_4,176_ = 1.1, *p* > 0.05). The rats were then divided into four groups: males EtOH_ND_, males EtOH_PD_, females EtOH_ND_, and females EtOH_PD_. Inactive lever responses remained very low and unaffected (data not shown).

**Figure 2.**
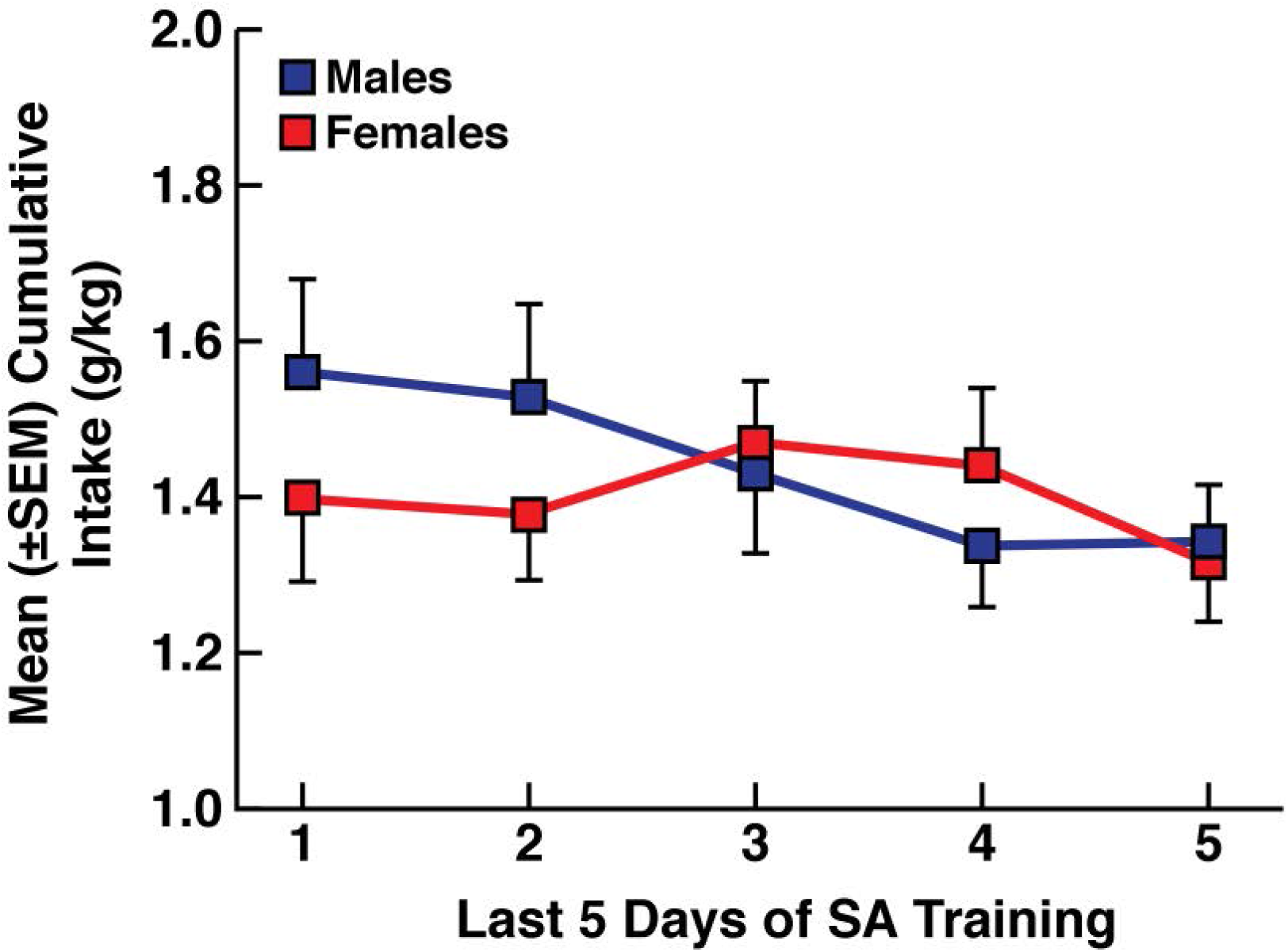
EtOH intake over the last 5 days of self-administration in daily 30 min sessions in males and females. *n* = 23 rats/group.

### EtOH self-administration during CIE exposure

Before CIE or air exposure, male and female rats were divided into two subgroups to obtain a similar baseline (BSL) of EtOH intake (males, *t*_21_ = 0.7, *p* > 0.05; females, *t*_21_ = 0.3, *p* > 0.05; Fig. 3A, B). EtOH intake during CIE exposure was not different between males and females. A difference in EtOH intake was observed between EtOH_ND_ and EtOH_PD_ at weeks 4, 5, and 6 of dependence induction (three-way ANOVA: sex, *F*_142_ = 2.8, *p* > 0.05; dependence, *F*_1,42_ = 49.0, *p* < 0.001; time [weeks 4, 5, and 6], *F*_3,126_ = 11.2, *p* < 0.001; sex × dependence × time interaction, *F*_3,126_ = 0.4, *p* > 0.05; Fig. 3A, B). A significant increase in EtOH intake was observed in EtOH_PD_ subjects at weeks 4, 5, and 6 compared with EtOH_ND_ (two-way ANOVA in males: group [EtOH_ND_ *vs*. EtOH_PD_], *F*_1,21_ = 36.2, *p* < 0.001; time [baseline, week 4, week 5, and week 6], *F*_3,63_ = 4.9, *p* < 0.01; group × time interaction, *F*_3,63_ = 19.7, *p* < 0.001; two-way ANOVA in females: group [EtOH_ND_ *vs*. EtOH_PD_], *F*_1,21_ = 18.7, *p* < 0.05; time [baseline, week 4, week 5, and week 6], *F*_3,63_ = 7.0, *p* < 0.001; group × time interaction, *F*_3,63_ = 4.6, *p* < 0.01). Compared with the training (BSL) condition, EtOHPD subjects exhibited a significant increase in EtOH intake from week 4 to 6 (*p* < 0.001, Tukey *post hoc* test following one-way repeated-measures ANOVA; males: *F*_10,30_ = 13.2, *p* < 0.001; females: *F*_10,30_ = 6.4, *p* < 0.05). No differences were observed between EtOHND and the training condition. Inactive lever responses remained very low and unaffected (data not shown).

**Figure 3.**
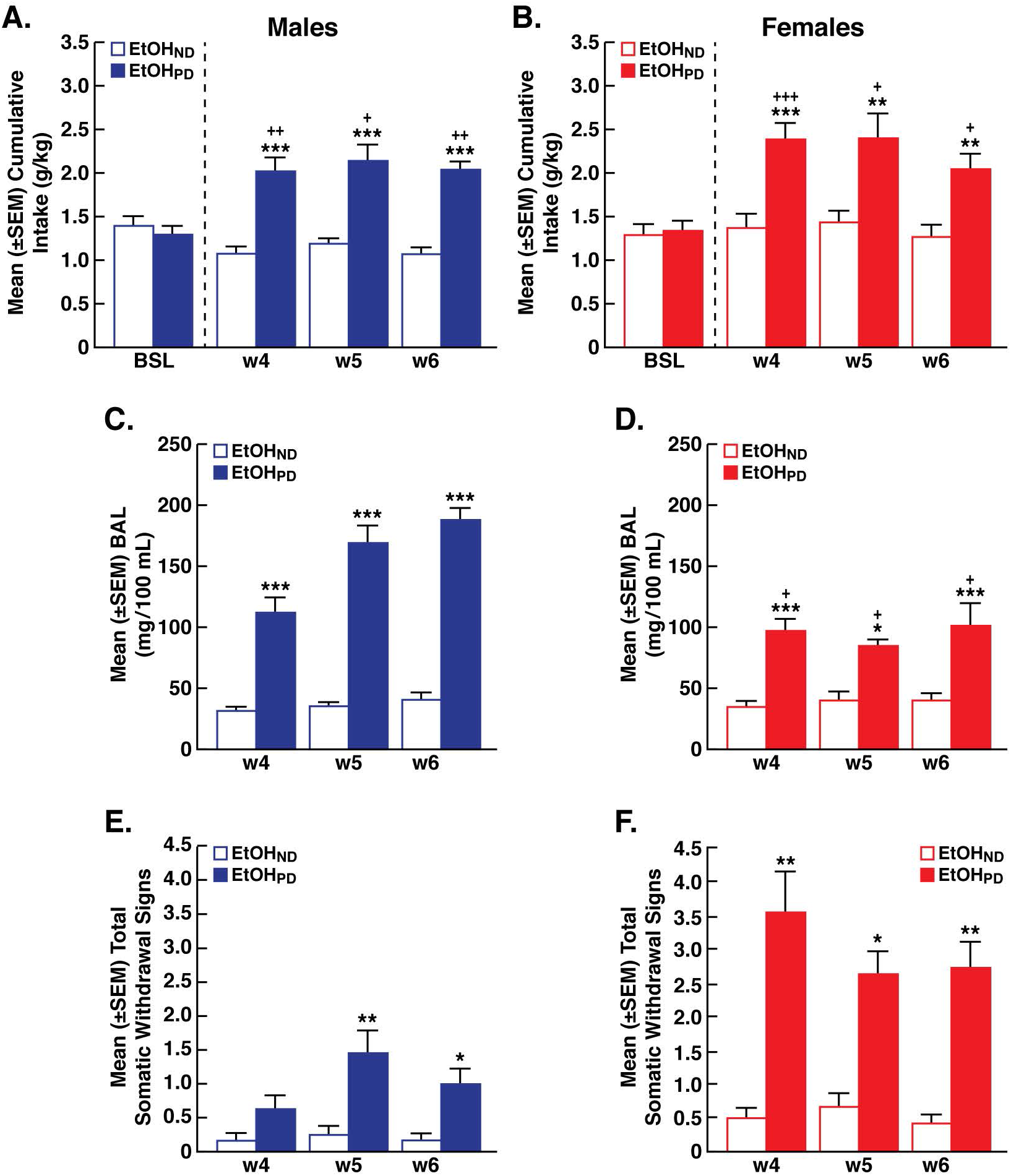
Cumulative EtOH intake was higher in EtOH_PD_ animals compared with EtOH_ND_ animals and compared with the training condition in both males (**A**) and females (**B**). Blood alcohol levels (BALs) that were measured immediately after the self-administration session during CIE vapor exposure were higher in EtOH_PD_ animals compared with EtOH_ND_ animals in males (**C**) and females (**D**). Somatic withdrawal signs (WDS) that were recorded prior to EtOH self-administration sessions during CIE vapor exposure at 8 h abstinence were higher in EtOH_PD_ animals than in EtOH_ND_ animals in males (**E**) and females (**F**). ^+++^*p* < 0.001, *vs*. respective BLS (Tukey *post hoc* test); \**p* < 0.05, \*\**p* < 0.01, \*\*\**p* < 0.001, *vs*. EtOH_ND_ (Tukey *post hoc* test). *n* = 11/12 animals/group. Basal level of EtOH intake during training (BSL).

### Blood alcohol levels

Blood alcohol levels were measured after each self-administration session, which were higher in EtOHPD animals compared with EtOHND animals during weeks 4, 5, and 6 of CIE exposure (*p* < 0.001, Tukey *post hoc* test following one-way ANOVA; males: *F*_5,63_ = 43.5, *p* < 0.001; females: *F*_5,63_ = 11.20, *p* < 0.001; Fig. 3C, D).

### Withdrawal scoring

During weeks 4, 5, and 6 of CIE vapor exposure, somatic withdrawal signs that were measured before each self-administration session were higher in EtOH_PD_ animals than in EtOH_ND_ animals (males: *p* > 0.05 at week 4, *p* < 0.01 at week 5, and *p* < 0.05 at week 6, Dunn’s *post hoc* test following Kruskal-Wallis *H* = 22.7, *p* < 0.001, Fig. 3E; females: *p* < 0.01 at week 4, *p* < 0.05 at week 5, and *p* < 0.01 at week 6, Dunn’s *post hoc* test following Kruskal-Wallis *H* = 37.8, *p* < 0.001, Fig. 3F).

### Effects of NTX on EtOH self-administration during abstinence

#### EtOH intake

Separate analyses of the differential behavioral effects of NTX were performed between males and females.

In males (Fig. 4), NTX reduced EtOH intake at all abstinence time points in EtOH_ND_ animals, whereas a trend toward a reduction appeared at L-Abst and was significant at P-Abst in EtOH_PD_ animals (three-way ANOVA: dependence, *F*_1,19_ = 23.7, *p* < 0.001; NTX treatment, *F*_1,19_ = 15.9, *p* < 0.001; abstinence time point, *F*_2.38_ = 15.9, *p* < 0.001; dependence × NTX treatment × abstinence point interaction, *F*_2,38_ = 1.9, *p* > 0.05; Fig. 4A-C). Naltrexone decreased EtOH intake in EtOH_ND_ rats at all abstinence time points (A-Abst: *t*_10_ = 7.5, *p* < 0.001; L-Abst: *t*_10_ = 2.3, *p* < 0.05; P-Abst: *t*_10_ = 5.5, *p* < 0.001; Fig. 4A-C), whereas the effect of NTX in EtOH_PD_ rats was significant only at P-Abst (A-Abst: *t*_9_ = 0.7, *p* > 0.05; L-Abst: *t*_9_ = 2.1, *p* > 0.05; P-Abst: *t*_9_ = 4.1, *p* < 0.01; Fig. 4A-C). The analysis of the percent intake (relative to vehicle) following NTX treatment (Fig. 4, inset) confirmed that NTX produced differential effects between EtOH_PD_ and EtOH_ND_ rats at A-Abst (Bonferroni *post hoc* test: *p* < 0.01, *vs*. EtOH_PD_ following two-way ANOVA: group [EtOH_ND_ *vs*. EtOH_PD_], *F*_1,10_ = 2.477, *p* > 0.05; time [A-Abst, L-Abst, P-Abst], *F*_2,20_ = 10.70, *p* < 0.001; group × time interaction: *F*_2,20_ = 6.727, *p* < 0.01).

**Figure 4.**
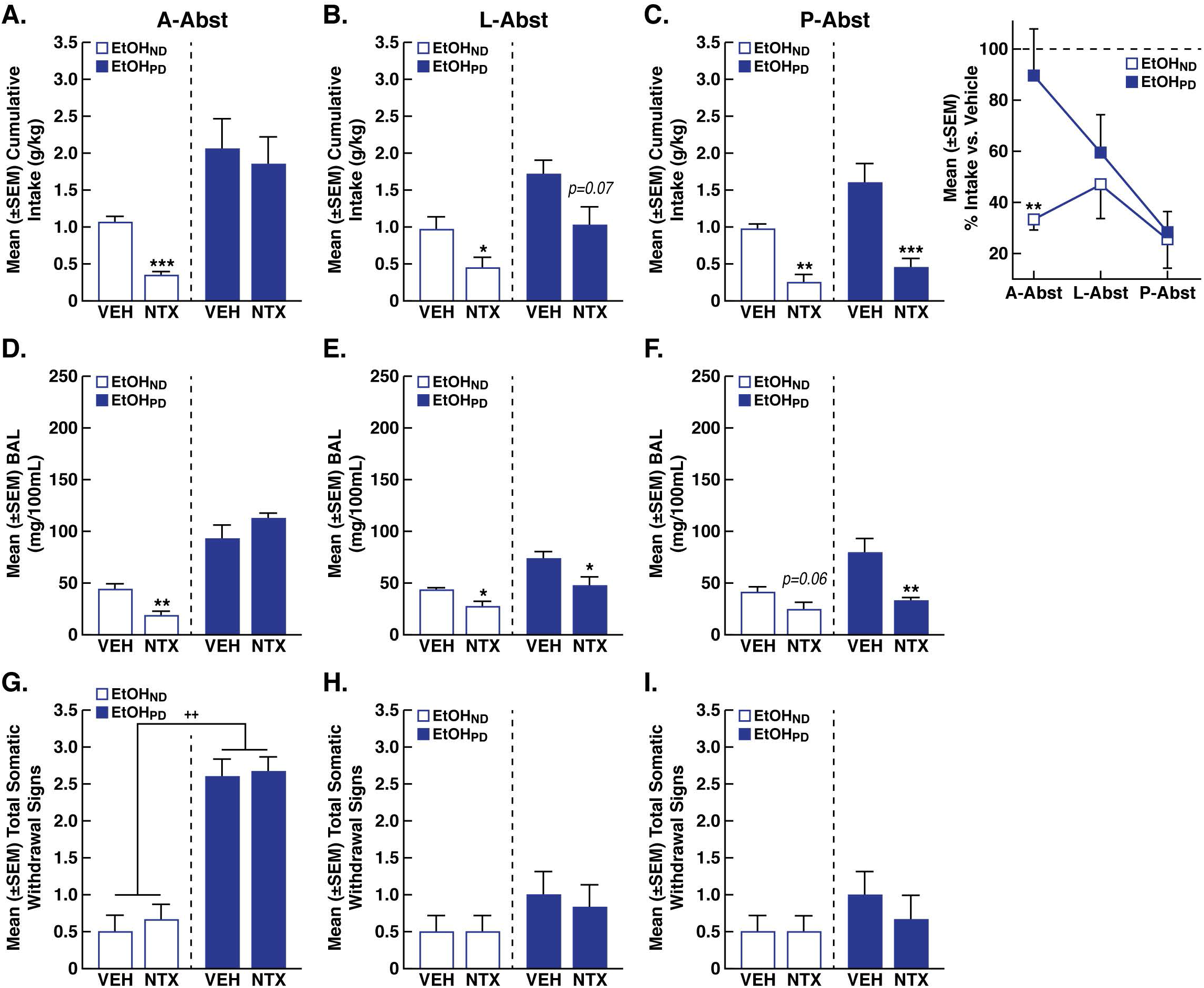
Effects of NTX in males. Cumulative EtOH intake after NTX or VEH administration in EtOH_PD_ animals *vs*. EtOH_ND_ animals at A-Abst (**A**), L-Abst (**B**), and P-Abst (**C**). Blood alcohol levels (BALs) were measured immediately after self-administration following NTX or vehicle (VEH) administration in EtOH_PD_ and EtOH_ND_ animals at A-Abst (**D**), L-Abst (**E**), and P-Abst (**F**). Somatic withdrawal signs (WDS) were monitored prior to EtOH self-administration following NTX or VEH administration in EtOH_PD_ and EtOH_ND_ animals at A-Abst (**G**), L-Abst (**H**), and P-Abst (**I**). \**p* < 0.05, \*\**p* < 0.01, \*\*\**p* < 0.001, *vs*. respective VEH; ^++^*p* < 0.01, *vs*. EtOH_ND_. *n* = 5/6 animals/group. (**Inset**) Percent EtOH intake following NTX relative to VEH administration at A-Abst, L-Abst, and P-Abst. *p < 0.05, vs. EtOHPD. *n* = 5/6 animals/group.

In females (Fig. 5), NTX reduced EtOH self-administration in both EtOH_PD_ and EtOH_ND_ rats similarly at the three abstinence time points (three-way ANOVA: dependence, *F*_1,19_ = 11.0, *p* < 0.01; NTX treatment, *F*_1,19_ = 18.9, *p* = 0.001; abstinence time point, *F*_2,38_ = 1.4, *p* > 0.05; dependence × NTX treatment × abstinence time point interaction, *F*_2,38_ = 0.3, *p* > 0.05; Fig. 5A-C). Separate *t*-tests confirmed that NTX reduced EtOH intake in both EtOH_ND_ and EtOH_PD_ rats at all abstinence time points (EtOH_ND_: A-Abst, *t*_10_ = 2.2, *p* < 0.05; L-Abst, *t*_10_ = 2.5, *p* < 0.05; P-Abst, *t*_10_ = 2.9, *p* < 0.05; EtOH_PD_: A-Abst, *t*_9_ = 2.5, *p* < 0.05; L-Abst, *t*_9_ = 4.8, *p* < 0.001; P-Abst, *t*_9_ = 3.4, *p* < 0.01; Fig. 5A-C). The analysis of percent intake (relative to vehicle) following NTX treatment (Fig. 5, inset) confirmed that EtOH_PD_ and EtOH_ND_ females responded similarly to NTX treatment at all abstinence time points (two-way ANOVA: group [EtOH_ND_ *vs*. EtOH_PD_], *F*_1,10_ = 0.3, *p* > 0.05; time [A-Abst, L-Abst, P-Abst], *F*_2,20_ = 0.2, *p* > 0.05; group × time interaction, *F*_2,20_ = 0.2, *p* > 0.05; Fig. 5, inset). Inactive lever responses remained very low and unaffected (data not shown).

**Figure 5.**
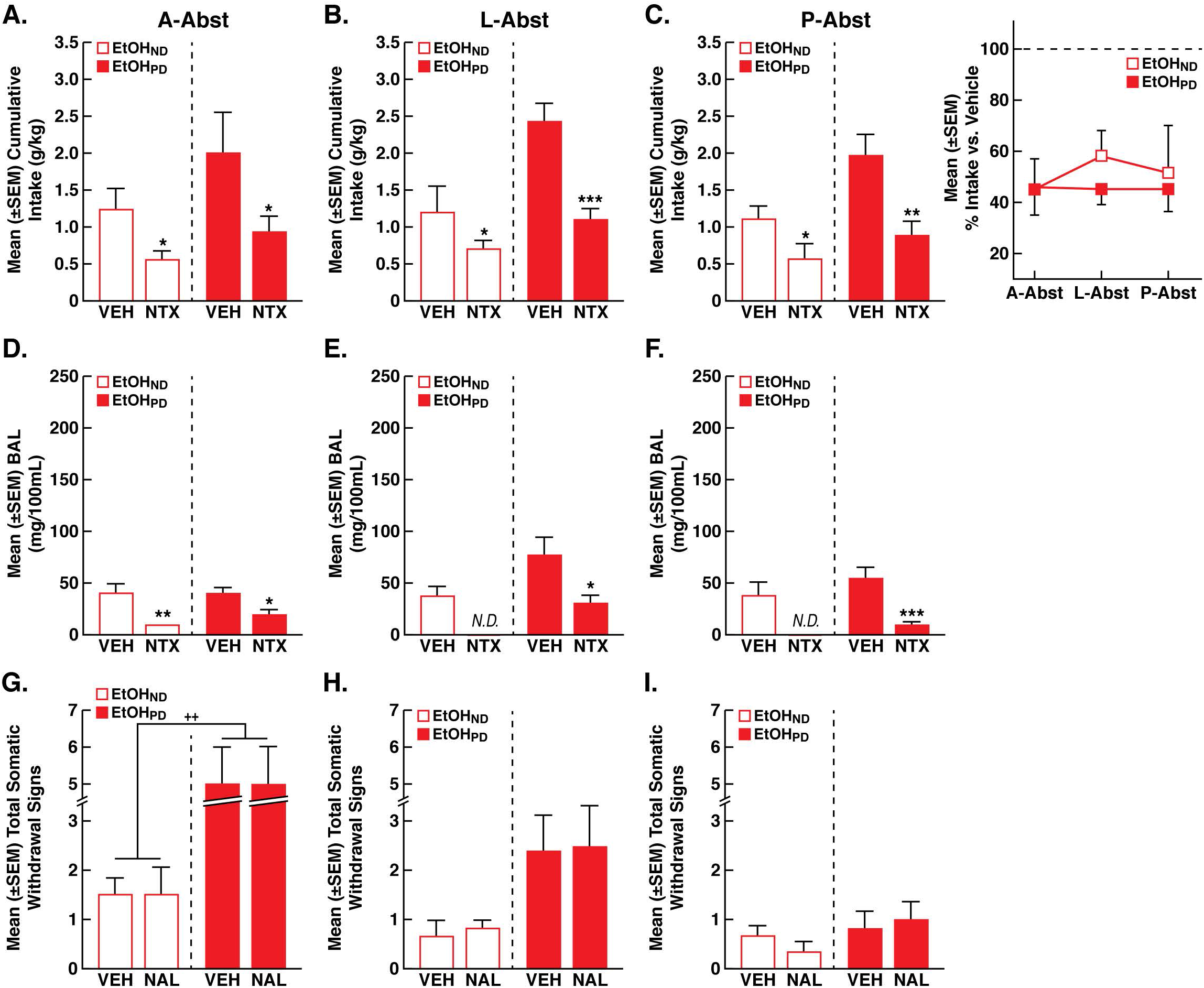
Effects of NTX in females. Cumulative EtOH intake following NTX or vehicle (VEH) administration in EtOH_PD_ animals *vs*. EtOH_ND_ animals at A-Abst (**A**), L-Abst (**B**), and P-Abst (**C**). Blood alcohol levels (BALs) were measured immediately after self-administration following NTX or VEH administration in EtOH_PD_ and EtOH_ND_ animals at A-Abst (**D**), L-Abst (**E**), and P-Abst (**F**). ND, not detectable. Somatic withdrawal signs (WDS) were recorded immediately before self-administration following NTX or VEH administration in EtOH_PD_ and EtOH_ND_ animals at A-Abst (**G**), L-Abst (**H**), and P-Abst (**I**). \**p* < 0.05, \*\**p* < 0.01, \*\*\**p* < 0.001, *vs*. respective VEH; ^++^*p* < 0.01, *vs*. EtOH_ND_. *n* = 5/6 animals/group. (**Inset**) Percent EtOH intake following NTX *vs*. VEH administration at A-Abst, L-Abst, and P-Abst. *n* = 5/6 animals/group.

#### Blood alcohol levels

In EtOH_ND_ males, the NTX-induced decrease in EtOH self-administration was associated by a significant lower BAL at A-Abst and L-Abst in EtOH_ND_ rats (A-Abst: *t*_10_ = 3.6, *p* < 0.01; L-Abst: *t*_10_ = 2.6, *p* < 0.05; P-Abst: *t*_10_ = 2.6, *p* = 0.06; Fig. 4D-F). In EtOH_PD_ rats, lower BALs in NTX-treated animals *vs*. vehicle-treated animals was observed at L-Abst and P-Abst but not at A-Abst (A-Abst: *t*_9_ = 1.5, *p* > 0.05; L-Abst: *t*_9_ = 2.3, *p* < 0.05; P-Abst: *t*_9_ = 3.5, *p* < 0.01; Fig. 4D-F).

In females, the NTX-induced decrease in EtOH intake in both EtOH_PD_ and EtOH_ND_ animals was associated with a significant lower BAL (*vs*. vehicle treatment) at A-Abst, L-Abst, and P-Abst (EtOH_ND_: A-Abst, *t*_10_ = 3.8, *p* < 0.01; BALs were under the detection limit in NTX-treated animals at L-Abst and P-Abst; Fig. 5D-F; EtOH_PD_: A-Abst, *t*_9_ = 2.5, *p* < 0.05; L-Abst, *t*_9_ = 2.5, *p* < 0.05; P-Abst, *t*_9_ = 4.8, *p* < 0.01; Fig. 5D-F).

#### Somatic withdrawal signs

Somatic withdrawal signs that were measured before each self-administration session (Fig. 4G-I, 5G-I) were higher in EtOH_PD_ rats compared with EtOH_ND_ rats at A-Abst in both males (Dunn’s *post hoc* test following Kruskal-Wallis *H* = 17.8, *p* < 0.001) and females (Dunn’s *post hoc* test following one-way Kruskal-Wallis *H* = 12.1, *p* < 0.01), whereas no difference in somatic withdrawal signs were found between EtOH_ND_ and EtOH_PD_ at L-Abst (Kruskal-Wallis: males, *H* = 0.7, *p* > 0.05; females, *H* = 7.2, *p* > 0.05) and P-Abst (Kruskal-Wallis: males, *H* = 0.6, *p* > 0.05; females, *H* = 2.5, *p* > 0.05).

## Discussion

One characteristic of clinical AUD is that dependent subjects will consume EtOH to relieve or avoid withdrawal symptoms (in rats, this usually starts to occur after 6-8 h of abstinence; Peer et al., 2013). The present study strongly supports the validity of the CIE vapor exposure model as a valid procedure to induce and study EtOH dependence in both male and female rats. Indeed, EtOH_PD_ rats exhibited an enhancement (i.e., escalation) of EtOH drinking during dependence (week 4 to week 6), suggesting a transition from controlled to excessive EtOH intake most likely in an effort to alleviate negative withdrawal states (for review, see Koob, 2014). These behavioral changes are characteristic of dependence and reflect neuroadaptive changes that are induced by chronic intermittent EtOH that in turn disrupts brain function (e.g., reward), physiological processes, and cognitive processes (Koob, 2008; Becker and Mulholland, 2014) and could account for changes in the efficacy of NTX to reduce EtOH drinking in the present study.

No differences in EtOH intake were observed between males and females in the present study during training and CIE vapor exposure. This was associated with similar BALs and somatic withdrawal signs in both sexes (Fig. 3). The lack of a difference between males and females is difficult to reconcile with previous studies that reported a range of contradictory results. Several studies have demonstrated sex differences in EtOH intake, in which female rats drank more than males (Blanchard et al., 1993; Morales et al., 2015; Walker et al., 2008; Li and Lumeng, 1984), whereas other studies found no differences (van Haaren and Anderson, 1994; Moore and Lynch, 2015; Schramm-Sapyta et al., 2014). The use of different strains (e.g., Wistar, Sprague-Dawley, and Long Evans) or different experimental designs (e.g., two-bottle choice, self-administration, and CIE vapor exposure) may account for such discrepancies. For example, female Long Evans and Wistar rats consumed more EtOH than their male counterparts when given either continuous or intermittent access to EtOH in their home cages (Priddy et al., 2017). Under operant conditions, no sex or strain differences were found in drinking prior to the development of EtOH dependence (Priddy et al., 2017). Consistent with the present results, upon dependence induction by CIE exposure, Wistar rats of both sexes substantially escalated their EtOH intake compared with their nondependent drinking levels, whereas Long Evans rats only exhibited a moderate escalation of drinking, without showing any sex difference (Priddy et al., 2017). Thus, strain, sex, and drinking condition may interact to modulate EtOH drinking and are important factors to consider when exploring individual differences in EtOH drinking and dependence.

The neurobiological basis for sex differences in AUD is largely unknown, partially because most studies of EtOH drinking are conducted in males only. However, sex differences are currently considered clinically (Agabio et al., 2016). The present results showed that both male and female Wistar rats acquired EtOH self-administration and consumed the same amount of EtOH through the training phase. During the final 3 weeks of the CIE procedure (week 4 to week 6), both males and females exhibited an equivalent and significant escalation of EtOH intake compared with their matching air-exposed control groups. However, males and females responded differently to NTX. In EtOH_ND_ animals (both males and females), NTX significantly reduced EtOH intake at all stages of abstinence (A-Abst, L-Abst, and P-Abst), and a significant change in the efficacy of NTX was observed in EtOH_PD_. Naltrexone was effective in EtOH_PD_ males only at P-Abst but reduced EtOH drinking in EtOH_PD_ females at all three abstinence time points.

Nondependent males and nondependent and postdependent females presented a NTX-induced decrease in EtOH drinking, regardless of the duration of abstinence. However, postdependent males became responsive to NTX over time and after a longer period of abstinence. The change in the efficacy of NTX is consistent with earlier findings that showed that NTX suppressed EtOH consumption in nondependent subjects (Gonzales and Weiss, 1998; Coonfield et al., 2002; Shoemaker et al., 2002; Stromberg, 2004), and this effect was blunted in dependent subjects (Ji et al., 2008; Sabino et al., 2006; Sabino et al., 2013; Walker and Koob, 2008). The efficacy of NTX in reducing EtOH intake may be affected by several factors, such as the model that is used (e.g., two-bottle choice *vs*. self-administration), the duration of EtOH self-administration, the severity of dependence, timing of the pharmacological intervention, sex, and changes in the endogenous opioid system. For example, acute NTX administration suppressed binge drinking in rats (Ji et al., 2008) and EtOH self-administration in dependent and nondependent animals during acute abstinence (Walker and Koob, 2008). Naltrexone was also highly effective in nondependent rats that were selectively bred for high alcohol preference (Sardinian alcohol-preferring [sP] rats; Sabino et al., 2006). Chronic NTX administration blocked the increase in EtOH consumption after a 5-day period of forced abstinence (Heyser et al., 2003). Changes in the endogenous opioid system are observed in response to chronic exposure to EtOH and after its removal, and these changes may be responsible for the effect of NTX or lack thereof. For example, chronic EtOH administration has been reported to increase activity of the hypothalamic β-endorphin system (Schulz et al., 1980; Angelogianni and Gianoulakis, 1993), cause no change (Seizinger et al., 1983), or even decrease activity (Scanlon et al., 1992). One hypothesis could be that the greater β-endorphin release that is induced by acute exposure to EtOH induces drinking, whereas a decrease in β-endorphin activity that is induced by chronic EtOH intake may promote and maintain EtOH consumption because of negative reinforcement rather than positive reinforcement. However, the reason why NTX was ineffective in postdependent males at A-Abst and L-Abst in the present study is not presently clear and will need to be explored further.

One tentative explanation for the differential effect of NTX between males and females may be differential changes in the endogenous opioid system in response to chronic EtOH. Overall, acute EtOH has been shown to stimulate the release of β-endorphin, enkephalins, and dynorphin in humans and rats (Gianoulakis et al., 1996a; Marinelli et al., 2003; Marinelli et al., 2005; Marinelli et al., 2006; Dai et al., 2005). Chronic EtOH exposure induced changes in opioid peptide systems that involve alterations at the levels of the peptides themselves (Gianoulakis et al., 1996b; Lindholm et al., 2000), changes in receptor densities and effector systems (Turchan et al., 1999; Chen and Lawrence, 2000), and modifications of mRNA that encode both opioid peptides and receptors (Przewlocka et al., 1997; Rosin et al., 1999). Therefore, the differential effects of NTX in males and females maybe reflect sexual dimorphism of the endogenous opioid system and different effects of chronic EtOH use in males and females. Sexual dimorphism in the endogenous opioid system has been described in the control of pain. For example, variable responses to the analgesic effects of opioids have been attributable to sex (Dahan et al., 2008), but the mechanisms that underlie sex differences in opioid analgesia remain elusive. Sex differences in opioid analgesia are also not likely related to differences in opioid receptor density because no differences between male and female rats in brain μ- or δ-opioid receptor populations have been found (Kepler et al., 1989). In humans, nonselective κ and μ ligands have been reported to produce stronger analgesic effects in women than in men (Rasakham and Liu-Chen, 2011). In animals, selective κ receptor agonists have been found to produce greater antinociceptive effects in males than in females (Rasakham and Liu-Chen, 2011). These observations suggest that the extent of sex differences in κ-, μ- and δ-mediated analgesia is related to species, strain, ligand, and pain model. For example, the selective κ-opioid receptor agonist spiradoline was more potent in female than in male Wistar rats in the tail-withdrawal test (Terner et al., 2003). The exact nature of the changes in the endogenous opioid system during EtOH dependence and withdrawal that can explain the higher sensitivity to NTX in EtOHPD females remains to be established.

The present results further support targeting the endogenous opioid system to prevent excessive drinking that is characteristic of AUD, even after long periods of abstinence. The present study extends our knowledge of the efficacy of NTX treatment by showing that it varies between sexes and different time points of abstinence. This could explain some of the contradictory findings that described the effects or lack of effects of NTX in some cases (Kakidani et al., 1982; Bell and Reisine, 1993). The present study has important implications because it points toward individual differences in the endogenous opioid system that should be considered when developing new pharmacotherapies to treat AUD.

## Acknowledgements

This is publication number 29664 from The Scripps Research Institute. The authors thank Maryjo Sampang for technical assistance and Michael Arends for assistance with manuscript preparation. The authors have no conflict of interest.

